# InstaPrism: an R package for fast implementation of BayesPrism

**DOI:** 10.1101/2023.03.07.531579

**Authors:** Mengying Hu, Maria Chikina

**Affiliations:** Department of Computational and Systems Biology, University of Pittsburgh

## Abstract

Computational cell-type deconvolution is an important analytic technique for modeling the compositional heterogeneity of bulk gene expression data. A conceptually new Bayesian approach to this problem, BayesPrism, has recently been proposed and has subsequently been shown to be superior in accuracy and robustness against model misspecifications by independent studies. However, given that BayesPrism relies on Gibbs sampling, it is orders of magnitude more computationally expensive than standard approaches. Here, we introduce the InstaPrism algorithm which re-implements BayesPrism in a derandomized framework by replacing the time-consuming Gibbs sampling steps in BayesPrism with a fixed-point algorithm. We demonstrate that the new algorithm is effectively equivalent to BayesPrism while providing a considerable speed advantage. InstaPrism is implemented as a standalone R package with C++ backend and can be accessed from GitHub at https://github.com/humengying0907/InstaPrism.

## Introduction

Bulk tissue transcriptome represents a mixture of gene expression signals from heterogeneous cell populations. The process of “deconvolution” aims to computationally separate these mixture signals and provide estimates of celltype abundance and in some cases estimate expression states of individual cell-type components. This problem is particularly of interest in cancer profiling as tumor samples are composed of malignant and non-malignant cells (the tumor microenvironment or TME). Understanding the interplay between malignant cells and the TME is an active area of research with therapeutic potential [1, 2, 3].

While the advent of single cell sequencing alleviates for computational deconvolution many studies cannot be performed with single cell assays exclusively due to resource constraints. Thus, the problem of deconvolution remains current, though many existing methods will assume that some matching single cell data is available [4, 5, 6, 7, 8].

Among numerous deconvolution methodologies that have been developed so far, BayesPrism [8] provides a novel Bayesian deconvolution framework that infers cell-type specific proportions and expression states using a reference scRNAseq dataset from similar tissue samples. Extensive analysis in the original BayePrism as well as independent deconvolution benchmarking studies [9, 10] have suggested that BayesPrism is superior to previous methods under different realistic settings. Specifically, BayesPrism appears highly robust to different sources of model misspecification that can introduce bias into standard reference-regression based methods, such as biological heterogeneity within cell-types [9] or technical differences between single cell reference and bulk [10])

However, due to its reliance on Gibbs sampling, BayesPrism shows a clear limitation in computational efficiency. For example, for a bulk expression matrix of 100 samples, BayesPrism needs hours of processing time [9] whereas competing methods take only seconds. The long running time limits the application of BayesPrism to large scale studies.

To address these limitations, we present InstaPrism, an R package that provides fast implementation of BayesPrism. Maintaining the same conceptual framework and corresponding generative model, InstaPrism replaces the time-consuming Gibbs sampling steps with a fixed-point algorithm. It produces nearly identical deconvolution results while reducing the running time to seconds, making it possible to utilize the BayesPrism approach in large-scale analysis.

## Package Overivew

### InstaPrism Input

InstaPrism has exactly the same input as BayesPrism, namely bulk RNAseq and annotated single-cell RNAseq and for a comparable tissue type. The single cell data is expected to be annotated to cell-type and cell-state within a specific cell-type. In the original paper, cell-states for malignant cells correspond to cells from a single patient. Cell-states for other cell-types can be specified according to single cell clustering. This cell-state specification is critical for achieving robustness against biological heterogeneity.

InstaPrism also accepts a BayesPrism constructed `Prism` object as input, which can be built via standard frameworks in the BayesPrism package.

### The InstaPrism Algorithm

We consider a single bulk gene expression sample with *G* genes, *X*_*G×*1_, and a cell-type/cell-state reference matrix, *A*_*G×S*_, where *S* is the total number of cell-states. It is important to note that both *X* and *A* are specified in terms of read counts. BayesPrism infers two variables, the proportions of cell-states in *X, θ*_*S×*1_ and another variable *Z*_*G×S*_. *Z* represents the allocation of the read counts in *X* to different cell-types. In the InstaPrism algorithm we keep track of another matrix called *B*_*G×S*_ which depends on both the static reference input *A* and the current proportion estimate *θ*. Each row of *B* is the mean of a multinomial distributions for assigning the reads of gene *g∈* [1, *G*] to the *S* different cell-states. This is a reformulation of the two-step multinomial model described in BayesPrism. In particular, at each iteration we maintain ∑_*g*_ *B*_*g,s*_ = 1 for all *g∈* [1, *G*] by normalizing the rows.

Overall the parameters of InstaPrism can be summarized as follows:

- Input: *X*_*G×*1_: gene expression in a given bulk sample in counts
- Input: *A*_*G×S*_: gene expression cell-states in counts
- Output: *Z*_*G×S*_: cell-state specific gene expression in counts
- Output: *θ*_*S×*1_: cell-state fractions
- Intermediate variable: *B*_*G×S*_: probability of assigning a read in gene *g ∈* [1, *G*] to a cell-state *s ∈* [1, *S*]

Given these variable definitions, InstaPrism can be described as the following deterministic algorithm. We use notation *Z*_*g*,_ and *Z*_,*s*_ to indicate genes/rows and cell-states/columns of *Z* respectively.

#### Algorithm 1 Fixed-point algorithm

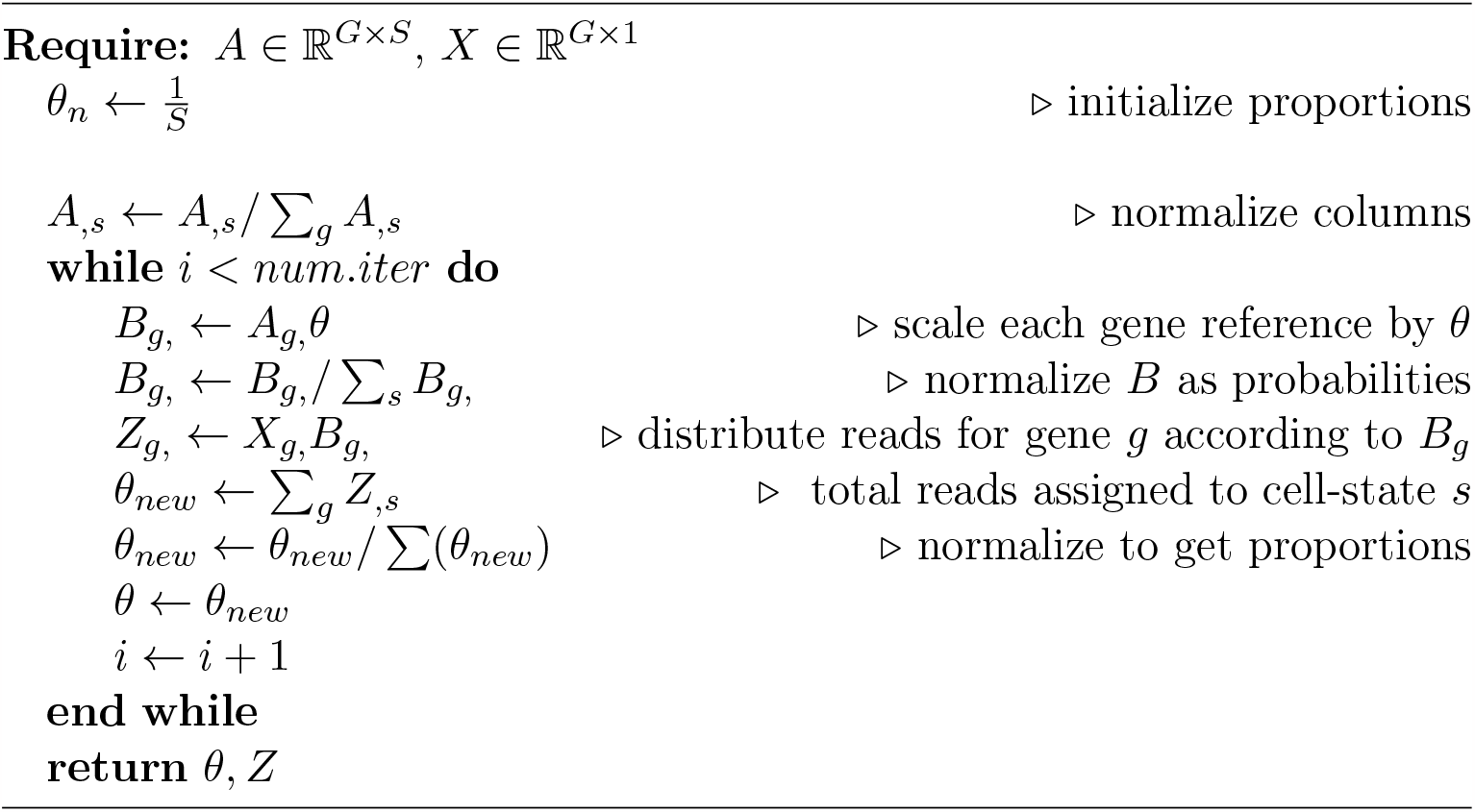

The InstaPrism algorithm essentially finds *θ*, which is the fixed point of the above iterative updates. The fixed point *θ* has the property that assigning each gene’s read counts according to renormalized *A*_*i*,_*θ* and forming the per cell-state deconvolved matrix *Z* obeys the equality ∑_*g*_ *Z*_,*s*_ = *Cθ* (where *C* is the total counts in *Z*).

The algorithm can also be seen as a derandomization of the original BayesPrism method. Where BayesPrism samples from Dirichlet and Multinomial distributions InstaPrism simply takes the expectation. We also combine two sequential multinomial steps described in BayesPrism equations 4 and 5 into one multinomial allocation per gene with mean parameters described by *B*. By avoiding the sampling the running time is significantly improved.

### Output

Under default settings, InstaPrism returns cellular fraction estimates at celltype level as well as cell-state level for each bulk RNAseq sample. InstaPrism can also output the cell-state and/or cell-type specific gene expression for each sample in the bulk dataset which is a three-dimensional expression array. InstaPrism also supports the original `updateReference` module from BayesPrism and will optionally return the updated cell-type fraction estimates. However, we find that this step changes the results very little but incurs additional computational cost.

### Comparison with BayesPrism

Using the tutorial data provided in the original BayesPrism package, which consists of both single-cell reference and bulk RNA-seq to deconvolve, we compared the deconvolution results from InstaPrism and BayesPrism. We find that the cell-type fraction estimates from two methods are nearly identical. This is true using either scRNAseq-based reference (Fig. 1) or the updated reference.

**Figure 1:**
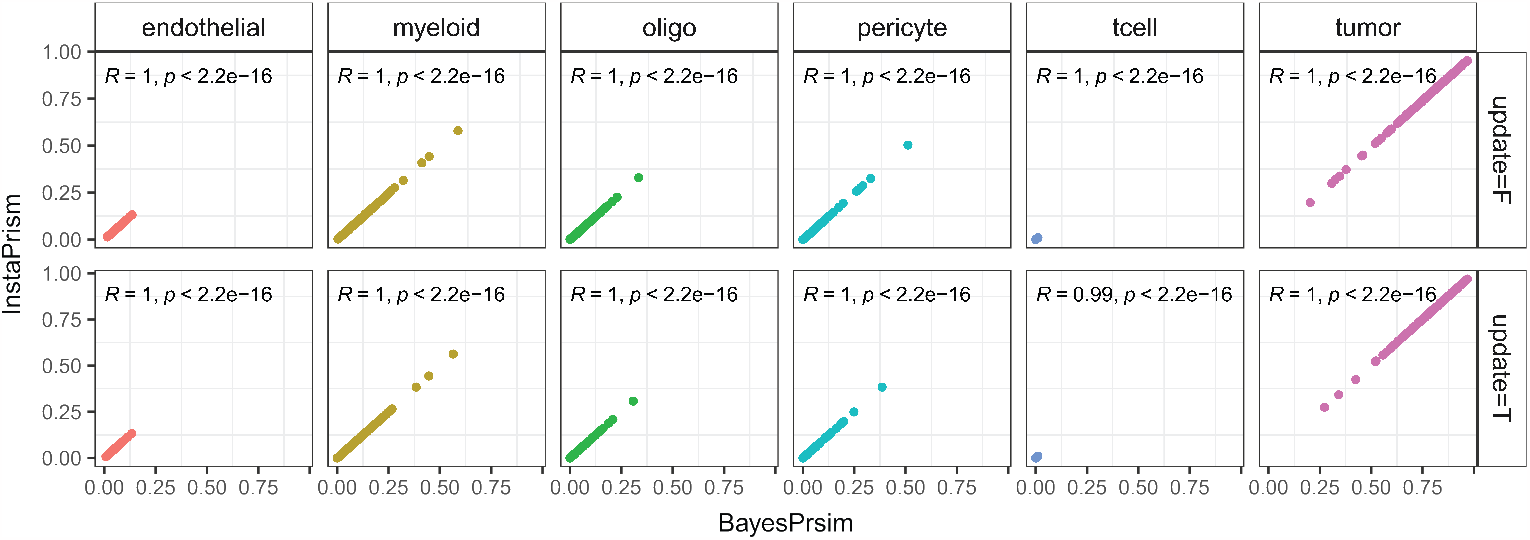
Deconvolution results comparison. Scatter plot comparing cell.type fraction estimates in 169 bulk samples from InstaPrism and BayesPrism, using either initial sc-RNA based reference or updated reference.

Both InstaPrism and BayesPrism update the cell fraction over iterations. Comparing how the fraction estimates change over iterations, we find that two methods followed the same update trajectory (Fig. 2) further supporting our assertion that InstaPrism is equivalent to BayesPrism minus the sampling. We note however, that as expected in the absence of sampling the update trajectories are smoother.

**Figure 2:**
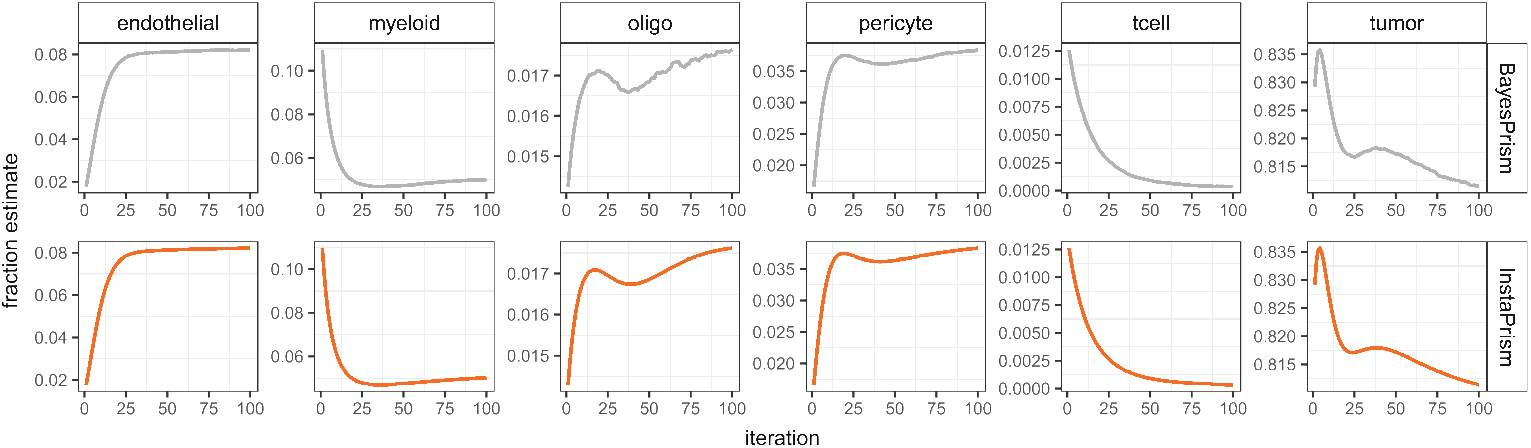
Fraction update comparison. Lineplot showing how cell-type fraction change over iterations in BayesPrism and InstarPrism for one bulk sample.

### Running time

InstaPrism greatly reduced running time by replacing the time-consuming Gibbs sampling step with a fixed-point algorithm. Below we showed how running time differs during the update steps in Fig 2.

At each iteration, InstaPrism takes only around 1/65 the running time of BayesPrism (Fig 3a). Importantly, since each InstaPrism iteration already effectively computes an expectation, we are able to reduce the total number of iterations. The dataset used in (Fig 1 and 2), which has 73 unique cell-states, we set number of iterations equal to 100, as we found that fraction estimates after 100 iterations show little difference (with absolute difference less than 0.01). To be conservative we set the default number of iterations to either 20 or 2 times the total number of cell-states, whichever is greatest. The parameter can also be set by the user.

**Figure 3:**
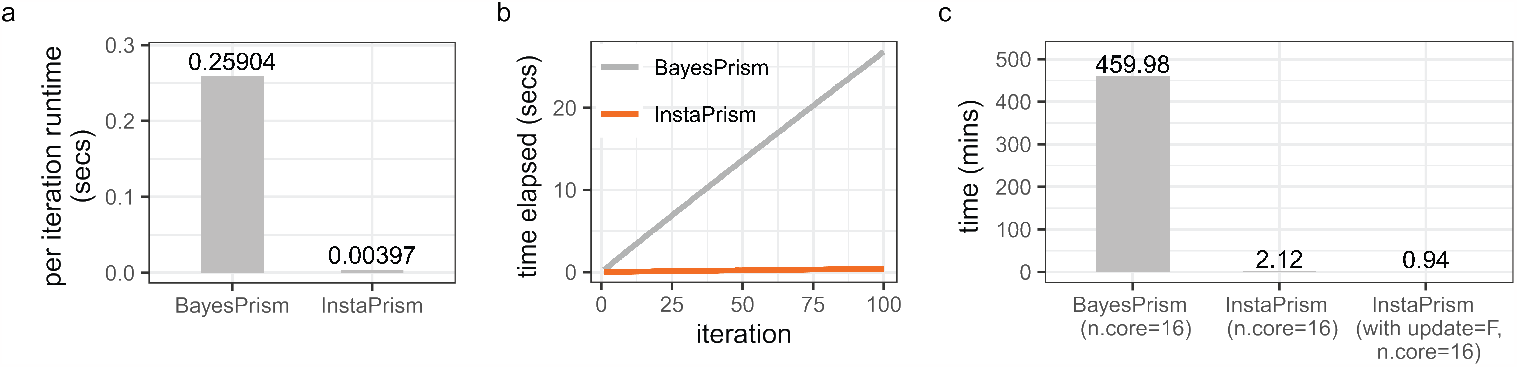
Running time comparison. (a) Barplot showing average runtime needed to update fraction in a single iteration for one bulk sample. (b) Line-plot showing total time elapsed over 100 iterations when updating a single cell.state fractions in one bulk sample. (c) Barplot showing total time required to get deconvolution results for the tutorial data provided in BayesPrism.

We reported the total deconvolution time of BayesPrism and InstaPrism on the tutorial data provided in the BayesPrism package (Fig 3c). InstaPrism significantly accelerated the deconvolution process and reduced the running time to only seconds under the default deconvolution procedure (with `update=F` argument).

## Conclusion and discussion

InstaPrim provides a fast implementation of the Bayesian-based deconvolution model. It takes advantage of the conceptual Bayesian framework proposed in BayesPrism but reformulates it as closed-form updates. The deconvolution results of InstaPrism is highly comparable with BayesPrism and can be implemented for large-scale deconvolution study. The improved running time will also enable exploration of how different cell-state and reference specifications affect the results.

The simplified algorithm also clearly showcases the conceptual difference and advantage of the BayesPrism framework. Standard reference/regression-based methods, such as CibersortX [11], can be viewed as modeling the *expected* gene expression given cell-type proportions. These methods differ in terms of reference construction and the loss used to fit the regression problem. On the other hand, the BayesPrism framework treats the data itself as a hard constraint. It is also notable that the reference matrix is essentially renormalized to probabilities and as such only the relative expression of a gene across cell-types is taken into account. It is likely that this feature is responsible for the notable robustness property of BayesPrism.

## Availability

The R package of InstaPrism can be accessed from GitHub at https://github.com/humengying0907/InstaPrism.

